# Entrainment of network activity by closed-loop microstimulation in healthy ambulatory rats

**DOI:** 10.1101/2020.07.10.196725

**Authors:** Alberto Averna, Page Hayley, Maxwell D Murphy, Jimmy Nguyen, Stefano Buccelli, Federico Barban, Randolph J. Nudo, Michela Chiappalone, David J. Guggenmos

## Abstract

As our understanding of how motor output is generated increases, it is clear that there is a need to understand the interactions of multiple distinct regions rather than just the output properties of primary motor cortex. This becomes even more imperative when trying to understand how different regions may contribute to recovery following injury. In this study we used a technique that promotes functional motor recovery after injury, activity-dependent stimulation (ADS), to determine the short- and long-term effects on network activity and neuroplasticity of intracortical connections. ADS uses recorded neural activity to trigger stimulation of the brain and may be utilized to manipulate neuronal connectivity *in vivo*, representing a novel technique to shape intrinsic neuroplasticity. The aim of this work was to compare the effect of ADS to randomly-generated stimulation (RS) of the somatosensory area (S1) on the single units’ patterns of activity taking place in the premotor cortex (RFA) and to investigate whether synaptic plasticity changes occur in S1 as a consequence of 21 consecutive days of stimulation. In particular, we examined both firing rate changes and correlation between spiking activity and stimuli in chronically-implanted healthy ambulatory rats during both spontaneous and evoked activity, resulting from the two stimulation paradigms. Finally, we evaluated changes in expression of synaptophysin at the end of the treatment. This experimental procedure demonstrated the ability of ADS to modulate firing properties of RFA within daily recording sessions and to promote synaptogenesis in S1, further strengthening the idea that this Hebbian-inspired protocol can be used to effectively modulate cortical connectivity and thus suggesting its translational potential for promoting recovery after brain injury.

## Introduction

The brain is composed of networks comprised of by both anatomically and functionally connected regions that support sensory and motor function (Levy *et al*., 2012). For instance, the primary motor cortex is not solely responsible for motor output, but it is actively involved in the processing of somatosensation, thanks to its anatomical and functional connections with the somatosensory cortex, thalamus and premotor cortices (Romo *et al*., 1993; Nudo *et al*., 1996; Bolognini *et al*., 2016). Following injury or disease, disconnection can occur, due to the disruption of communication between coordinated regions. This can lead to sensorimotor or cognitive dysfunctions described as ‘disconnection syndromes’ (Catani & ffytche, 2005; Bullmore & Sporns, 2009), that are associated with several neurological and neuropsychiatric conditions, such as schizophrenia (Micheloyannis *et al*., 2006; Rubinov *et al*., 2009; Friston *et al*., 2016; Kottaram *et al*., 2019), Alzheimer disease (Sorg *et al*., 2007; Supekar *et al*., 2008; Badhwar *et al*., 2017; Marchitelli *et al*., 2018) and autism spectrum disorder (Turner *et al*., 2006; Just *et al*., 2012; Hernandez *et al*., 2015; Catani *et al*., 2016). However, these impairments can also result from direct tissue loss, as in case of stroke (Wang *et al*., 2010; Grefkes & Fink, 2011; Carter *et al*., 2012; Adhikari *et al*., 2017) and traumatic brain injury (TBI) (Nakamura *et al*., 2009; Messé *et al*., 2013; Sharp *et al*., 2014; Caeyenberghs *et al*., 2017). Impairment from stroke, one of the most common causes of adult onset disability, stems from a combination of direct tissue loss and the disruption of sensorimotor integration leading to changes in motor performance (Machado *et al*., 2010). Rehabilitation from these sensorimotor deficits relies on re-establishing functional connectivity, including sensorimotor integration.

Non-invasive techniques to assess functional connectivity between brain regions such as fMRI, have been important in understanding large scale alterations in patterns of activity following injury (Cramer & Bastings, 2000; van Meer *et al*., 2010; Rehme *et al*., 2011; Rehme & Grefkes, 2013). Unfortunately, as with many non-invasive techniques, there is a trade-off in both spatial and temporal resolution, and these techniques are susceptible to distortion and other artifacts. Conversely, neural signals recorded invasively can provide a much more detailed picture of activity, opening avenues for looking at very specific alterations in communication in disconnection syndromes and in novel therapies for treating these disconnections (Engel *et al*., 2005).

While physical therapy is the standard-of-care for recovering lost function following injury, its effects are often limited or incomplete. Restoring function following impairment to improve quality of life is a primary challenge in scientific and clinical research. Translational approaches attempt to address this need through innovative invasive therapeutic approaches. Recent studies (Adkins-Miur, 2003; Kleim *et al*., 2003; Jackson *et al*., 2006; Guggenmos *et al*., 2013) demonstrated that electrical stimulation can be used successfully to manipulate neuronal functional connectivity, representing a potentially powerful tool to assist in guiding appropriate *de novo* functional connections to restore lost sensorimotor integration. While artificially building functional neural connections following injury is the ultimate goal, there are a number of more basic, mechanistic questions on how these connections can be shaped *in vivo*, especially in healthy, intact brains. Previously, we demonstrated that ADS of the primary somatosensory cortex (S1) is able to alter the evoked response of the Rostral Forelimb Area (RFA, the premotor cortex equivalent in the rat) by modulating cortico-cortical connectivity in healthy anaesthetized rats within a single recording session (Averna *et al*., 2019), but questions of how this stimulation impacts the connectivity longitudinally in non-anesthetized rodents remain. To this end, we investigated the effects of two different intracortical microstimulation paradigms to S1 on the firing properties of neural populations within the pre-motor cortex of the intact rat. Activity-Dependent Stimulation (ADS), is a technique for artificially pairing the activity of two populations of neurons by coupling the activity of one neuron (the ‘Trigger’ neuron) to stimuli delivered at a separate location. Prolonged ADS has shown success in promoting behavioral recovery in rodents following focal injury of primary motor cortex (Guggenmos *et al*., 2013), but there is little information on how chronic ADS may impact firing in normal animals. Performing these protocols on healthy rodents will allow us to determine the effect of the electrical stimulation on cortico-cortical pathways.

In the present study we investigated the ability of ADS to alter the normal cortical firing patterns under a condition in which it is unlikely that *de novo* axodendritic sprouting would occur. We acquired neurophysiological signals from RFA of healthy, ambulatory rats during stimulation of S1 using either ADS or random stimulation (RS). Our recordings consisted of 200-minute long sessions recorded over the course of 21 consecutive days from rats allowed to freely move within a testing chamber. We found that ADS was effective for both entraining the network response by facilitating stimulus associated neuronal activity in the motor areas and promoting synaptogenesis in the sensory cortex, compared with RS. Importantly, our results strengthen the feasibility of using Hebbian inspired stimulation approaches in novel therapies for patients living with disabilities such as stroke, TBI, or other conditions in which neural communication is disrupted.

## Methods

### Animals and dataset

All experiments were approved by the University of Kansas Medical Center Institutional Animal Care and Use Committee. In total, 12 adult male Long-Evans rats (weight: 350-400 g, age: 4-5 months; Charles River Laboratories, Wilmington, MA, USA) were used in this study (Table 1).

**Table 1:**
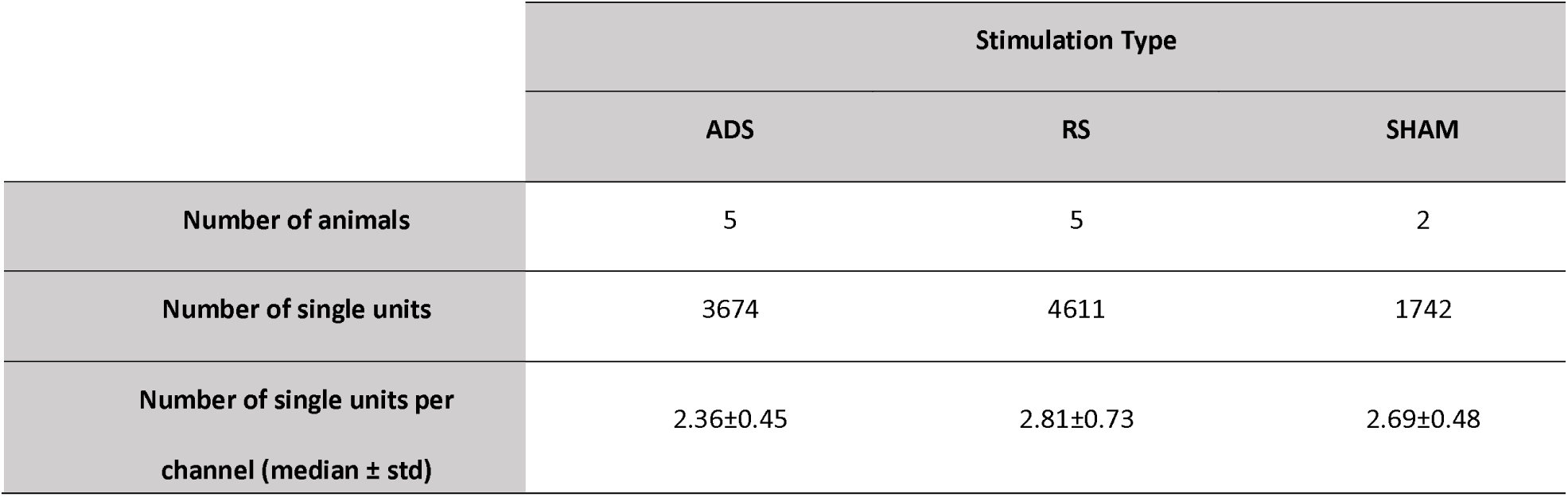
Healthy Awake dataset. Grouping of Rats into Activity-Dependent Stimulation (ADS), Random Stimulation (RS) and Control (SHAM).

|  | Stimulation Type |  |  |
| --- | --- | --- | --- |
|  | ADS | RS | SHAM |
| Number of animals | 5 | 5 | 2 |
| Number of single units | 3674 | 4611 | 1742 |
| Number of single units per channel (median ± std) | 2.36±0.45 | 2.81±0.73 | 2.69±0.48 |

### Surgical Procedures

Anesthesia was induced with gaseous isoflurane prior to surgery within a sealed vaporizer chamber, followed by injections of ketamine (80-100 mg/kg IP) and xylazine (5-10 mg/kg). Maintenance boluses of ketamine (10-100 mg/kg/h ip or im) were repeatedly injected as needed throughout the procedure. A stereotaxic frame was used to secure the rat heads, and an anal temperature probe was used to monitor the rats’ temperature. The rats’ eyes were protected with ophthalmic ointment. Either Lidocaine/Prilocaine cream or bupivacaine was applied to the scalp prior to performing a skin incision spanning rostro-caudally between ∼6 mm rostral to bregma and ∼5 mm distal to the atlanto-occipital junction. The cisterna magna or upper vertebrae were exposed by reflecting the overlying neck muscles. The spinal dura located in the foramen magnum was slightly punctured to control brain edema by allowing cerebrospinal fluid (CSF) drainage. After the retraction of the temporalis muscle, a craniectomy was performed to expose the primary motor (Caudal Forelimb Area: CFA) and premotor (RFA) cortical areas. The electrophysiological procedures were facilitated by the removal of the dura and application of sterile silicone oil to the cortex. Following the craniotomy, six holes were created using a small drill bit for the skull screws and anchoring rod. The skull screws were implanted into the parietal and intraparietal bones and secured with dental acrylic. An additional dose of Penicillin (15,000U) was injected subcutaneously at the end of the surgical procedure to prevent bacterial infections.

### Mapping Cortical Areas

The location of the RFA was determined using standard intracortical microstimulation (ICMS) protocols (Kleim, 2003). Upon completion of the craniectomy and durectomy, a picture of the vascular pattern of the cortical surface was taken and uploaded to a graphics program (Canvas GFX, Inc., Plantation, FL, USA), and a 250 µm grid was overlaid onto the image. At grid intersections, a glass microelectrode (10-25 µm diameter) filled with saline was inserted into the cortex at a depth of ∼1700 µm. Stimulus trains composed of 13 pulses lasting 200 µs each at 333 Hz were applied at 1 Hz intervals using a stimulus isolator (BAK Electronics, Umatilla FL, USA). The current was increased until visible movement around a joint was observed, up to 80 µA. RFA was defined by forelimb motor responses bordered caudally by neck and trunk responses and medially by face and jaw movements (Kleim *et al*., 1998). The forelimb sensory fields in S1 were identified with multi-unit recordings. A single-shank Michigan style electrode with sixteen recording sites (NeuroNexus, Ann Arbor, MI) was advanced into the sensory cortex to span across cortical layers II-V. The evoked sensory activity was monitored through an audio speaker during the procedure. Further characterization of the spikes was performed by amplifying, digitizing, filtering, and displaying the recorded neural activity on a display screen (TDT, Alchulta, FL). The electrode penetration sites that responded to palpation in the digits, paw and wrist were defined as forelimb sensory fields.

### Experimental Protocol

A four-shank, sixteen-contact site electrode with 1-1.5 MΩ impedance at each site (A4×4-3mm-100-125-177-CM16LP, NeuroNexus) was chronically implanted into the RFA at a maximum depth of 1600 μm (Figure 1A). The probe was coupled to an active unity gain connector that was connected to the recording system (TDT, Alchulta, FL) through an amplifier. All channels of neural data were sorted on-line using Principal-Component Analysis (PCA) trained on the first five minutes of recording sessions (TDT, Alchulta, FL). A single contact site that detected the neural spiking data at a moderate rate of spontaneous activity (4-10 Hz) was selected as the trigger channel for stimulating a cortical site in the forelimb-responsive S1.

**Figure 1.**
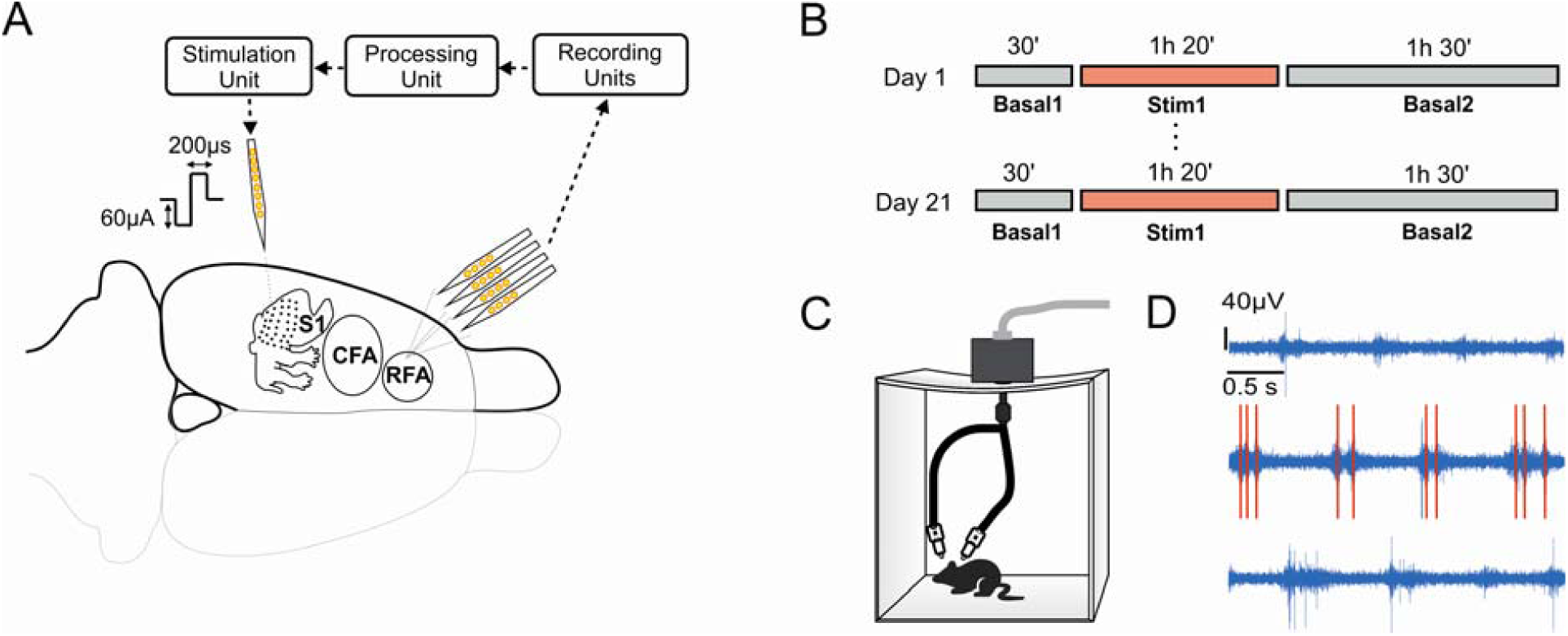
Experimental scheme of an implanted rat. A) Two multisite electrodes were implanted in the left hemisphere; one spike from a single channel in the L-RFA was used to trigger stimulation in the somatosensory area (in the case of ADS). B) Timeline of the protocol. Each rat was recorded for 21 consecutive days with each session containing two recordings of basal activity (30 min and 1 h and 30 min) and one stimulation session (1 h and 20 min) in between. C) Rats could freely move inside a transparent cage during the experimental session. D) A sample trace recorded during the three experimental phases (red lines represent the deleted stimulation artifacts).

A second sixteen-contact electrode (A4×4-3mm-100-125-703-CM16LP, NeuroNexus) was chronically implanted into S1 and used for stimulation through a single contact with ∼200 KΩ impedance (Figure 1A). A stimulus isolator and passive headstage (MS16 Stimulus Isolator, TDT) delivered the stimulus. Each neuronal spike profile recorded from a single contact in the RFA triggered a single 60 μA biphasic, cathodal-leading stimulus pulse (200 μs positive and 200 μs negative) in S1 after a 10 ms delay. In the RS designated rats, randomized stimuli (Poisson distribution centered approximately at 7 Hz) that approximated the observed frequency in the ADS rats (4-10 Hz) were used. Each stimulus was followed by a 28 ms blanking period to prevent runaway feedback from stimulus-evoked RFA spikes and stimulus artifacts from re-triggering stimulation (Figure 1D). Rats were allowed to recover five days post-operatively before initiating the study. All groups were recorded daily for 21 days. Each animal could move freely inside a self-made plastic cage (50×50×60 cm) for the entire duration of each experimental session but was not required to perform any sensorimotor tasks (Figure 1C). The daily recording was divided into the set *E* ={*Stim,Pre,Post*}, enumerated as follows: an 80-minute period of stimulation (‘Stim’) flanked by a 30-minute period (before stimulation, ‘Pre’) and a 90-minute period (after stimulation, ‘Post’) of no stimulation for a cumulative total of 3 hours and 20 minutes of recorded data per day (Figure 1B).

In the SHAM experiments, the daily recordings were performed identically, but the stimulation current was set to 0 μA for the entire 3 hours and 20 minutes of recording.

### Data Processing

Neural data was recorded (Tucker-Davis Technologies - TDT, Alchulta, FL, USA) and then processed offline. Streaming data was filtered through a band-pass filter (elliptic filter) in the bandwidth of the spikes (300-3000 Hz). A custom MATLAB (The Mathworks, Natick MA) script was used to identify the peaks in the extracellular electrical fields that corresponded to the action potentials of nearby neurons. Spikes in the filtered signal were detected using a combination of thresholds designed to robustly screen for unit activity as described previously (Bundy *et al*., 2019) Briefly, candidate spike samples were identified if the Smoothed Nonlinear Energy Operator (SNEO) (Mukhopadhyay & Ray, 1998) met or exceeded 4.5 times the median deviation of the SNEO time-series. The MATLAB (R2017a) ‘findpeaks’ algorithm was then used to identify peaks in the SNEO-valid reduced time-series, with the ‘MinPeakAmplitude’ parameter set to 4.5 times the median absolute deviation of the bandpass filtered time-series (Quiroga *et al*., 2004; Maccione *et al*., 2009)

After spikes were detected, a manual sorting stage was implemented to generate the set of all cluster indices *K*. During this stage, each voltage sample on the corresponding channel to the detected spike spanning 0.4-ms prior to 0.8-ms after each detected peak was used to perform a manual “cluster-cutting” (see the Matlab code at https://github.com/m053m716/CPLtools/tree/master/Matlab_Addons/Functions/Spike%20Analyses) procedure implemented in Matlab (R2017a) similar to existing packages such as MClust (Redish, 2008) or SimpleClust (Buccino *et al*., 2019). This custom interface provides a visual indicator of pairs of spike waveform features (e.g. the amplitude of two different time-samples relative to the peak) across the duration of the recording, such that cluster quality across the duration of a given recording was evident during the manual scoring procedure. Peaks that clearly corresponded to noise were excluded at this stage by assignment to a “noise” or “non-neural” source cluster (N_noise_ of N_Total_ spikes). Spikes were sorted for each recording separately. Clusters were not matched across days, therefore analyses did not require that cluster indices corresponded to the same source unit across days. In all subsequent analyses, the set of all detected, non-noise sorted spike peak time lists *P* was used to define subset *P*^*a,d*^(*k,e*) ∈ *P* of specific peak times detected for the index representing animal *a* ∈ *A*, on index *d* ∈ *D* the set of all post-operative days during epoch *e* ∈ *E* (i.e. *E* = {*Stim,Pre,Post*} for spike cluster index *k* ∈ *K*

### Quantification of relation between spiking (output) and stimuli (inputs): I/O Correlation

We tested the null hypothesis that RS and ADS invoke the same network response, as defined by the maximally correlated firing of the network (considered here as the “Output”) with the stimulus time-series (considered here as the “Input”) as already presented in (Scarsi *et al*., 2017). The correlation coefficient *R*(*X,Y,τ*) between two time-series processes *X* and *Y* depends on some offset lag *τ* such that sample *X* (*t)* is compared with sample *Y* (*t* − *τ)*. For discrete time-series, such as in this experiment, *τ* ∈ T(*L,w*) for *L* the maximum offset lag between the two time-series, and *w* the offset lag sample period (sometimes referred to as bin-width). Here, input ICMS pulse train *y*(*t*) ∈ *Y* and output spike train *x*(*t*) ∈ *X* were compared to find the maximum Input/Output (I/O) correlation for each epoch *e* ∈ *E* as *R*_*e*_(*X,Y,τ*_*max*_) by fixing *L* = 30*ms* and *w* = 1*ms* This short timescale was selected based on the primary interest, which was synaptogenesis resulting in de *novo* cortico-cortical synapses. Therefore, we selected *w* = 1*ms* due to the relatively high granularity of evoked output responses based on the estimated conduction velocity of corticospinal outputs (Stewart *et al*., 1990). We selected *L* = 30*ms* on the assumption that this should be a sufficient window to capture evoked monosynaptic spiking responses due to stimulation, based on an estimated cortico-cortical conduction velocity of between 0.10 m/s (Aroniadou & Keller, 1993) and 0.55 m/s (Murakoshi *et al*., 1993). Furthermore, these parameters match the timescale imposed previously in a hardware-constrained context (Guggenmos *et al*., 2013), allowing us to expand on those findings in the intact, ambulatory case.

We used the following algorithm to define each corrected I/O pairwise correlation 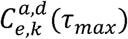, where *a* denotes the animal, *d* is the number of days after surgery, *e* is a member of *E* (the epochs), *k* is the spike cluster, and *τ*_*max*_ is the time lag for which the function 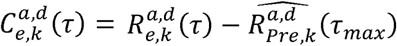is maximized. To compute the I/O pairwise correlation *R*_*e*_(*τ*)and the corresponding subtrahend 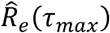,first, we defined each spike train 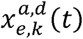, using 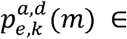 *P*, the time relative to the start of the sample record for the spike indexed by *m* 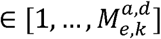 for 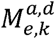spikes of unit *k* during epoch *e* on animal *a* and day *d*.

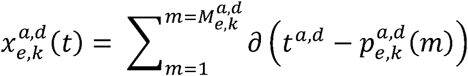

The Dirac delta function *∂(t*) is defined for all *t*^*a,d*^ time samples in the record as

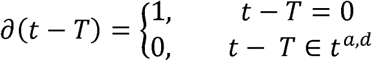

Similarly, we defined each matched stimulus train element 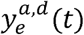, using 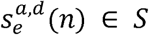, the time relative to the start of the sample record for stimulus indexed by 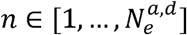 for ICMS pulses during epoch *e* on animal *a* and day *d*.

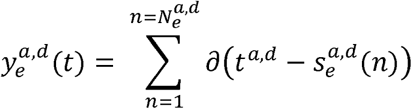

For notational convenience, we will drop the indexing scripts [*a,d,k*], assuming that any combinations from here on refer to index-matched elements unless an index is explicitly stated, and that any reference to *t* is specific to the experimental record from a single recording. To implement the convolution of *X* and *Y* used to recover *R*_*e*_*(τ*), a smoothing procedure was implemented in two steps: a binning of spike and stimulus times to yield 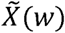 and 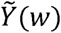, followed by a low-pass filtering step applied to each binned vector yielding 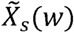 and 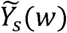.

This procedure could be implemented using the raw stimulus times as registered by the acquisition system when *e = Stim*; however, when *e* ∈ {*pre,post*}, by definition, ICMS pulses were not delivered. Therefore, for these epochs, two separate procedures *g* ∈ {*ADS,post*} were implemented for computing an estimate of the “sham” trigger I/O correlation 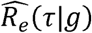In the case of *g* = *ADS*, we simply use the trigger unit times to act as a proxy for the stimulus train:

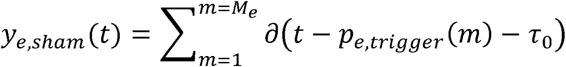

Where *τ*_0_ is the fixed offset latency of 10 ms between spike detection and stimulation when *e* = *stim.* However, for the case *g* = *RS* the choice of a “trigger” channel would be arbitrary. Because of the intrinsically higher expected correlation between ADS stimuli and observed spiking activity (*E*[*R* _*stim*_ (*x*_*trigger*_ (*t*),*Y* |*g* = *ADS, τ = τ*_0_)] = 1), no correction factor was subtracted for *g* = *RS* Therefore, the corrected I/O correlation is 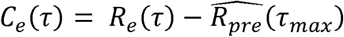, where

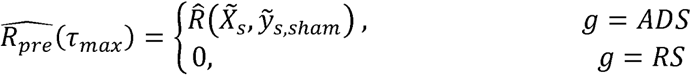

We also sought to distinguish the statistical effect peculiar to the observed patterns of stimuli, due to the fact that in general ICMS is known to exert alterations on extracellular activity. To that end, we generated 100 surrogate samples for each unit under consideration by randomly shuffling the observed spikes throughout each spike train, such that

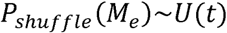

For the uniform random distribution over all observed sample times *U* (*t*), so that each shuffled surrogate list of spike times 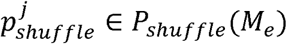 indexed by *j* ∈ [1,…,100] contains *M*_*e*_ spike times. These surrogate lists were then used to each compute 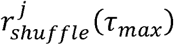 using the same steps as described previously. *C* (*τ*_*max*_) was considered statistically significant if it was greater than the 95^th^ percentile of the surrogate *R*_*shuffle*_ (*τ*_*max*_).

### Firing rate

Neuronal firing rates were evaluated before, during and after the stimulation by calculating the average firing rate during each period. Neurons whose firing rate was less than 0.01 spikes/s in the first basal period (Pre) were discarded. A bootstrapping method was used to determine if the difference in the firing rates of a given unit between two time-points significantly deviated from a null (zero centered) distribution (Slomowitz *et al*., 2015). The compared time segments (Pre, Stim and Post) were divided into 1-min bins, which were then randomly shuffled 10,000 times into two groups. The differences between the means of the randomly shuffled groups produced a null-distribution. The real difference was significant if it fell outside of the 95% confidence interval of the null-distribution.

### Post-Stimulus Time Histogram

The Post-Stimulus Time Histograms (PSTH) (Rieke *et al*.; Rieke, 1999) (1 ms bins, normalized across the total number of stimulation pulses) of the stimulus-associated action potentials of each sorted unit during the 28 ms following the stimulus pulses delivered from S1, has been calculated. The area under the normalized PSTH curve was used to quantify the total amount of stimulation-evoked neural activity during each stimulation phase. Only the units with significant responses were considered in this analysis (see Data Processing, I/O pairwise correlation).

### Immunochemistry

Following completion of all the recording sessions, rats were transcardially perfused with 4% paraformaldehyde and 0.1M phosphate buffered saline (PBS). Before histological processing, a fiducial marker was placed in the hemisphere contralateral to the implant along the lateral edge of the cortex. Coronal sections through the sensorimotor cortex were cut at 50μm kept in series with 300 μm between sections.

For processing, a blocking solution containing 5% normal goat serum (NGS) and 0.3% Triton X-100 in 0.1M PBS was pipetted on the tissue in each well and was rocked on a shaker table at room temperature for an hour. The solution was promptly removed, and the primary antibody solution was applied. The mouse-derived primary antibodies for synaptophysin (Millapore-Sigma) were diluted to 1:200 in 0.1 PBS with 2% NGS. 200μL of the solution was pipetted into each well and the trays were stored overnight at 4°C. The next day the sections were transferred into a new well tray with PBS and rocked for 10 minutes. This was repeated by changing out the PBS solution for a total of three rinse cycles. Following the final wash, the solution was pipetted off and 200μL of the secondary antibody solution was added to each well. The secondary solution contained a 1:200 dilution of goat anti-mouse IgGs with the conjugated fluorophore, Alexa Fluor^®^ 594, in 0.1M PBS. The secondary antibody solution was incubated for one hour and was washed with 0.1M PBS three times immediately after. The tissue sections were mounted on subbed slides and coverslipped using Hardset™ VECTASHIELD^®^ mounting medium with DAPI.

### Image Analysis

The slices were visualized using a Zeiss Axio Imager.M2 microscope. Representative sections within the rostral forelimb area and the sensory cortex, S1, were selected from each series by referencing an anatomical atlas and, when possible, the presence of electrode tracts to approximate electrode location (Paxinos *et al*., 2007). These slides were loaded onto the stage and the sections were digitally outlined at 25x magnification using the Stereo Investigator software (MBF Bioscience, Williston, VT, USA). The objective was switched to 20x (200x total magnification) and the slide was focused manually at pre-set sites across the tissue using the DAPI fluorescent channel. The light intensity was kept consistent at 20%. The exposure was set at 800ms for the GFP and DsRed channels and at 200ms for DAPI for all sections. Using the slide scanning workflow on Stereo Investigator, the camera took 200x images at each site using all three channels. These images were taken systematically across the tissue to produce a single tiled image of the coronal section for each channel.

Images were converted to 8-bit using the Fiji distribution of ImageJ (Schindelin *et al*., 2012). The images were then cropped to the area of interest on the superior aspect of the section. A bandpass filter was applied to remove the artifactual lines created in the image stitching process. The filter was designed using the FFT bandpass filter algorithm included in the Fiji process option, with the upper limit parameter set to 10,000 pixels while the bottom limit parameter was set to 1 pixel and the suppress stripes option enabled with a tolerance of 5%. A magenta look-up table was applied to the image and the histogram was adjusted to standardize the expression within the corpus callosum which has stable, low levels of synaptophysin. Regions of interest were created in Fiji for the left and right cortex. The cortical regions of interest (ROI) were defined by a straight line running parallel to the medial axis of the section that stretched from the medial to lateral edge of each hemisphere through the most superior point of the corpus callosum. The upper boundary of the ROI followed the superior border of the cortex.

A binary mask was generated in MATLAB (R2017a) using the ROIs traced in ImageJ. The mask was used to index a subset of pixels from each image, from which 500 random unique pixels were drawn from each hemisphere for a given section image. Each rescaled pixel intensity was also associated with the corresponding metadata pertaining to animal number (Rat); experimental group (Group: random stimulation, RS; or activity dependent stimulation, ADS); hemisphere (Hemisphere: left or right); and location (Area: rostral forelimb area, RFA; or sensory cortex, S1).The intensity, was fit using a Standard Least Squares model using the restricted maximum likelihood (REML) method in JMP 11 (SAS Institute Inc., Cary, NC). The model used the random effect of Rat, along with the full factorial combinations of the fixed effects of Group, Hemisphere, and Area, as well as a term to account for the Intercept.

## Results

### Daily changes in firing rate induced by ICMS

Changes in firing rate were evaluated over the entire duration of each recording session. The number of RS pulses was greater than the number of ADS pulses as well as the resulting mean firing rate observed during the stimulation phase (see Supplementary Materials, Figure S1 A, B). Since no changes were found across days (see Supplementary Figure S2) for each of the two stimulation paradigms (i.e. ADS or RS), different recording days were considered as different trials of the same animal and they were merged together for analysis (Figure 2A).

**Figure 2.**
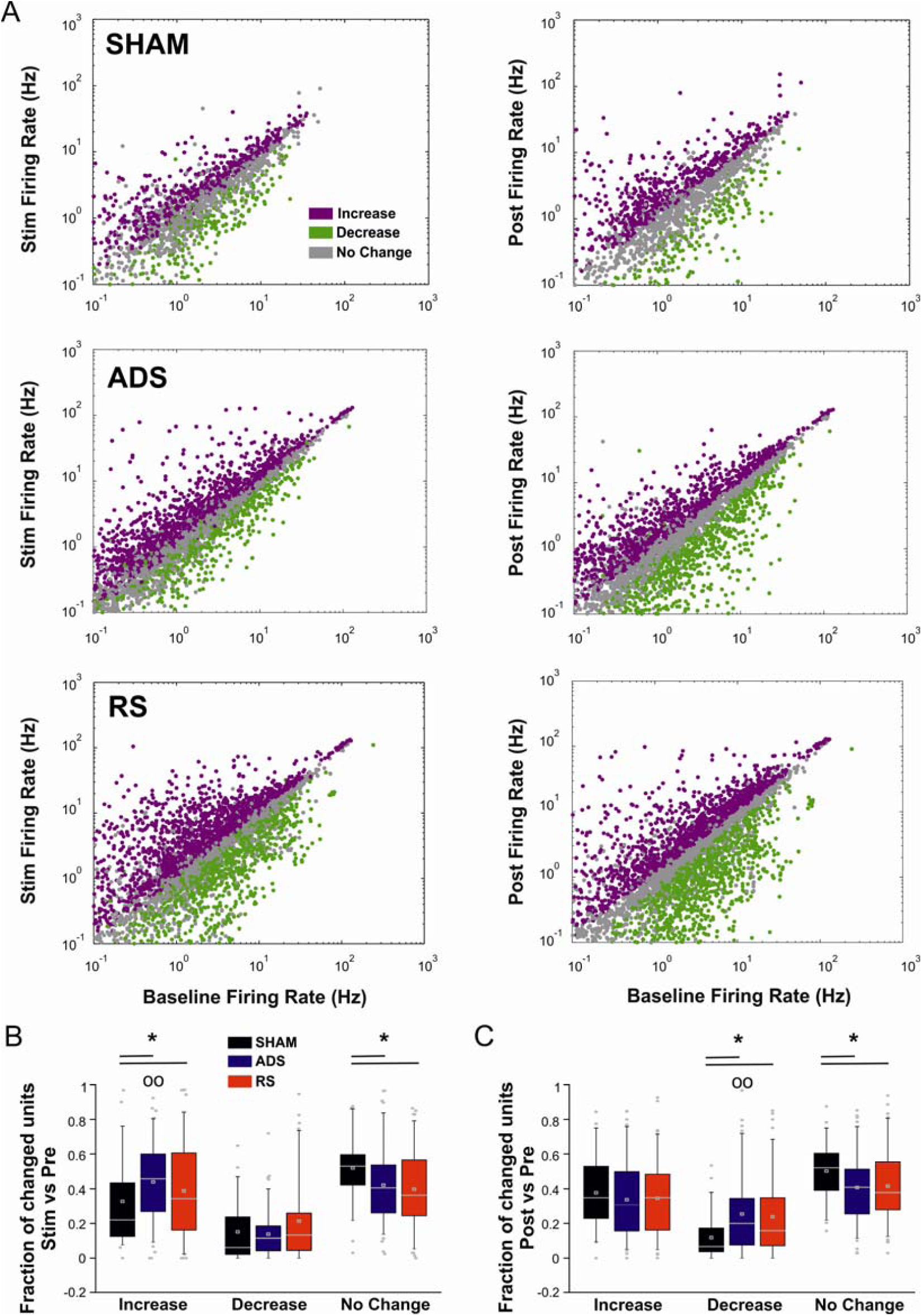
Effect of ICMS on Firing Rate. A) Per unit correlation between the baseline firing rates (x-axis, Baseline, i.e. 1Pre) and the firing rates during both the stimulation session (left, y-axis, Stim) and the Post stimulation session (right, y-axis, Basal post stimulation, i.e. Post) calculated per group (SHAM, ADS, and RS). Colors represent units that significantly increased (magenta), decreased (green) or remained stable (grey). B) Average fraction of units that significantly changed their firing during Stim with respect to the baseline period of recording (Pre) calculated in all three experimental groups. C) Average fraction of units that significantly changed their firing during Post with respect to the baseline period of recording (Pre) calculated in all three experimental groups (* ^O O^ p < 0.05; Kruskal-Wallis one-way analysis of variance of ranks with Dunnett’s multiple comparison test. ADS, RS vs SHAM within the same phase. p < 0.01; Kruskal-Wallis one-way analysis of variance of ranks with Dunnett’s multiple comparison test. Fraction of increased, decreased, no changed units versus different phases within groups (i.e. Pre vs Stim vs Post). For each box plot (B, C), the central white square indicates the mean, the central line illustrates the median and the box limits indicate the 25th and 75th percentiles. Whiskers represent the 5th and the 95th percentiles.

First, we calculated the number of units whose firing increased, decreased or remained constant between the stimulation phase and the baseline period of recordings (i.e. Pre). We found that for both ADS and RS paradigms, ICMS produced a significantly greater number of units showing increased firing rates compared to SHAM (Figure 2B). Furthermore, after the stimulation (i.e. during Post) the number of units which decreased their firing with respect to the baseline period was significantly higher for the stimulated animals than SHAM (see Figure 2C). These results were corroborated by the greater fraction of units that remained constant in the SHAM group in both Stim and Post (Figure 2B, C). Interestingly only for the ADS group, the fraction of increased units during Stim was significantly higher than during Post (within the same group), where the number of decreased units was greater than Stim (Figure 2B, C, ^OO^ marker).

### ADS produced more significant stimulus-associated responses than RS

The stimulus-evoked response was analyzed by post-stimulus time histograms and I/O pairwise correlation (cf. Data Processing in the Methods section). There was a blanking window of 4 ms following stimulus onset imposed to avoid false positive spikes from the amplifier coming out of saturation (Figure 3A). As shown in Figure 3B, the inter-stimulus interval distribution of the RS group was similar to that observed in the ADS group.

**Figure 3.**
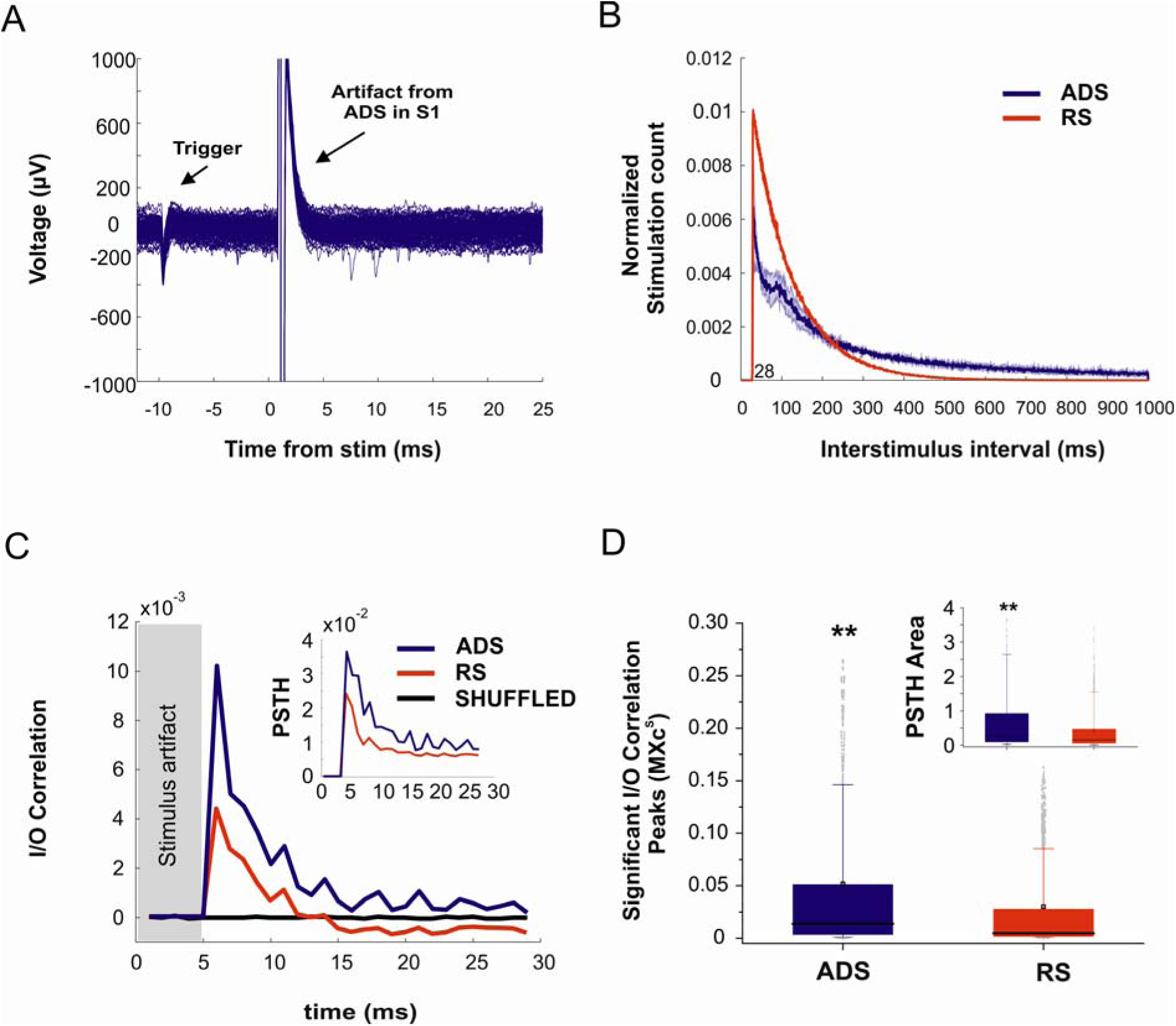
Evaluation of the stimulus-associated responses. A) Sample trace of recordings from the RFA showing stimulus artifacts from ADS delivered to S1 and activity of the trigger unit. In total, 100 superimposed traces are shown. B) Stimulation interval distribution (mean±STD, 1 s) in both representative ADS and RS subjects. Stimulation counts were normalized to the session length. C) I/O correlation traces (median) of the significant evoked responses in the two groups (ADS, blue and RS, red) calculated by pooling all 21 days of stimulation sessions. The median correlation trace for the shuffled trains is reported in black. Histograms traces portraying the average number of action potentials (median) discriminated from the neural recordings within 1-ms bins in 28 ms windows after each stimulation pulse in the two ICMS groups (ADS, blue and RS, red) is reported in the box (inset). D) Box Plots of the significant I/O correlation peaks in the two groups of ICMS. Box plots depicting the normalized PSTH areas (normalized to the total number of either ADS (Left) or RS (right) event) in the two groups of ICMS are reported in the box (inset). (**p < 0.001; Mann-Whitney Rank Sum test). For each box plot (D), the central black square indicates the mean, the central line illustrates the median and the box limits indicate the 25th and 75th percentiles. Whiskers represent the 5th and the 95th percentiles.

We evaluated the amount of spiking activity which was directly correlated to the stimulation pattern, delivered through one of the two protocols (i.e. ADS or RS). To consider the maximum correlations that were effectively induced by the stimulation, only the significant *C*_*e*_ (*τ*_*max*_) values with positive lags were considered in our analysis (see Data Processing, I/O pairwise correlation). Figure 3C shows the averaged (median) I/O correlation function of the significant evoked responses calculated for the two stimulation groups (ADS and RS). The correlation traces appear to be different for the two groups, with ADS exhibiting a greater level of stimulus-evoked activity than RS (Figure 3C, blue line). The PSTH derived from the neural recordings in the RFA by discriminating the spiking activity during the 28 ms after each S1 stimulus pulse, confirmed this result showing a higher number of spikes per stimulus for ADS (Figure 3C, top). Figure 3D shows the distribution of significant I/O correlation peaks. Even after the compensation procedure (see Data Processing), ADS peaks were significantly higher than RS (Figure 3D). Likewise, the stimulus evoked activity calculated in terms of PSTH area were greater for ADS (see Figure 3D, inset). No differences of both I/O correlation and PSTH area, were found across different days of recordings (see Supplementary Material, Figure S3).

### ADS increases Synaptophysin expression in S1

Following completion of treatment, we evaluated the expression of synaptophysin, a synaptic marker, for 11 subjects (i.e. 5 ADS, 5 RS and 1 SHAM). Visual inspection of cortical sections fluorescently stained for synaptophysin indicated a higher presence of synaptophysin in S1 near the electrode location (Figure 4A). To quantify this finding, a standard least-squares regression model was fit to predict synaptophysin expression using the fixed effects of treatment group (SHAM, ADS, or RS), region of interest (S1 or RFA), hemisphere (Left or Right), and their interaction effects, as well as the random effect of rat. The interaction of hemisphere and region (F ratio = 19.0012; p=<0.0001) was found to be significant, predicting greater fluorescent synaptophysin expression in the left (stimulated) S1 regardless of stimulation treatment. Furthermore, the interaction of group, hemisphere, and region was also significant (F ratio = 209.2172; p=<0.0001), indicating that ADS treatment predicted increased synaptophysin expression within the implanted left S1 over RS (Figure 4B). The poor linear model fit (RSquare = 0.108292) was expected due to the low overall synaptophysin expression as well as the relatively sparse presence of intense synaptophysin sources within each overall region of interest. There were no clear differences in synaptophysin expression within RFA across all groups.

**Figure 4.**
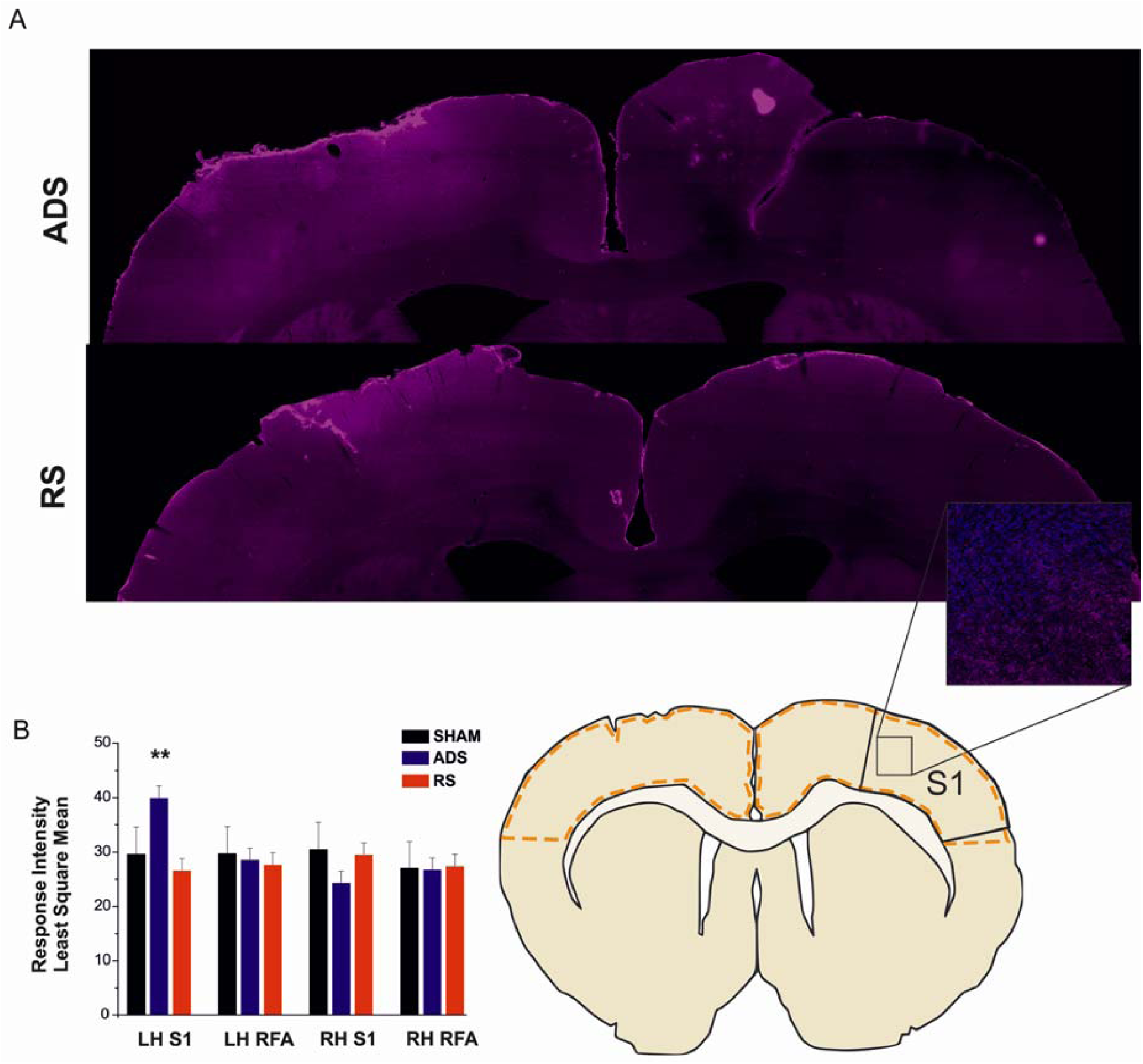
Fluorescent immunolabeling of synaptophysin expression in rat cortex after chronic electrode implantation. A) The panels show the representative S1 cortical area sections from each animal cropped and overlaid to highlight the comparative regions of interest. The top panel contains the right and left hemispheres of the ADS treatment side by side. Compared to the bottom panel which demonstrates the respective sections from the RS group, ADS treatment increases the synaptophysin expression with the implanted left S1. Also, of note is the consistency of synaptophysin expression in the un-implanted right hemisphere which is expected with normal expression as compared to the localized patterning in the left hemisphere—particularly with activity dependent stimulation. B) Synaptophysin expression levels in terms of Least Squares Means ± STD. **p<0.001, Friedman ANOVA with Dunn’s multiple comparisons. Inset is 200x magnification of an S1 representative section (based on (Paxinos & Watson, 2006)), where blue represents DAPI and magenta synaptophysin. Section outline shows the region of each hemisphere that was sampled for analysis. Inset shows expression of synaptophysin at the level of magnification used for analysis.

## Discussion

The purpose of this study was to determine whether it was possible to drive short- and long-term neurophysiological changes in response to cortical stimulation in healthy, ambulatory animals. We examined the response within pre-motor cortex (RFA) following stimulation within somatosensory cortex (S1) within individual recording sessions and any cumulative effect across multiple recording sessions. Further, we tested two types of stimulation, ADS (closed-loop activity-triggered stimulation) or RS (open-loop randomized stimulation), to see if synchronizing the activity between the regions impacted the degree to which we could observe changes in neuronal activity. Even if during the entire time of recording the spontaneous firing rate was not chronically modulated by the microstimulation, we found that both stimulation treatments induced daily changes in firing rate during the stimulation phase and the subsequent recording of spontaneous activity (i.e. Post) compared to SHAM. Moreover, ADS was more effective in entraining the network response by evoking stimulus-associated spiking activity than RS, whose evoked activity was less correlated with the stimulation train. Finally, we evaluated the amount of a synaptic plasticity marker (i.e. synaptophysin) present at the site of stimulation (S1) at the end of the 21 days experimental period. We found only ADS induced increased synaptophysin expression within the region of stimulation.

ICMS within the somatosensory cortex has been shown to be effective in altering the intracortical activity of the RFA in healthy anesthetized rats during a single recording session (Averna *et al*., 2019). ICMS, regardless of the type of stimulation (ADS or RS), induced an increase in the firing rate during a subsequent period of recording while ADS (but not RS) facilitated incremental increases in stimulus-associated activity over time, suggesting that this protocol may lead to a stronger association over more prolonged stimulation sessions. Although anesthetized preparations, as that described in our previous study, have numerous advantages since the state of the animal and the associated neurophysiological set-up can be considered stable over several hours, they also have several limitations. Ketamine anesthesia can have a significant impact on neuronal activity (Brown *et al*., 2010; Brown *et al*., 2011), influencing firing rate and synchrony in the hippocampus (Kuang *et al*., 2010) enhancing gamma oscillations acutely, but decreasing these same oscillations chronically (Ahnaou *et al*., 2017). Moreover, the modulation of GABAa receptor induced by ketamine produces opposite effects on different neuronal sub-population leading to a net cortical excitation in the prefrontal cortex (Homayoun & Moghaddam, 2007). The properties of inducing spiking changes with stimulation in a chronic preparation may be fundamentally different in a non-anesthetized preparation, especially when considering the role of spike-stimulation timing in ADS preparations. At the population level, the spatiotemporal structure of spikes firing in the awake cortex cannot be directly inferred from activity recorded in the anesthetized cortex, even when considering the same neuronal population (Greenberg *et al*., 2008).

Artificially pairing spike-firing in one populations of neurons with focal electrical stimulation of a second population of neurons can shape the efficacy of specific neural pathways *in vivo* (Jackson *et al*., 2006; Rebesco *et al*., 2010; Rebesco & Miller, 2011; Guggenmos *et al*., 2013; Nishimura *et al*., 2013). Temporal coding is critical for enhancing the network entrainment (synaptic efficacy) (Mainen & Sejnowski, 1995; Gal & Marom, 2013; Scarsi *et al*., 2017), and either ADS or Poisson distributed stimulation may similarly enhance the intrinsic pathways that would lead to increased firing rates in spontaneous activity.

In contrast to our previous findings in healthy anesthetized animals (Averna *et al*., 2019), we did not observe an increasing effect of ICMS (both ADS and RS) on firing rate. The fraction of neurons which increased their firing rate with respect to the baseline period (i.e. Pre) was not retained during Post (following stimulation), but we found a greater fraction of decreasing firing neurons compared to the non-stimulated controls (Figure 2C). These differences between the anesthetized and awake preparations may be explained by the different population-wide firing properties, such as the synchronization and relationship between the correlation and the firing rate, which are typically modulated by anesthesia (Greenberg *et al*., 2008).

During chronic recordings, we found ADS had a significantly higher proportion of neurons with increased firing rates during the Stim phase compared to the Post phase, but also a higher proportion of neurons that decreased firing rates from Post to Pre compared to the other stimulation conditions (Figure 2B, C). This would confirm that, although the response to ADS may differ between anesthetized and awake preparations, it is more effective at modulation firing activity in RFA than random stimulation conditions (Averna *et al*., 2019).

ADS is thought to induce changes in spiking probability through Hebbian mechanisms, therefore, a change in the post-stimulation activity in the trigger region should be observed (Jackson *et al*., 2006; Guggenmos *et al*., 2013; Averna *et al*., 2019). Indeed, ADS induced a greater level of stimulus-associated activity than RS and this evoked activity more correlated with the stimulation train (Figure 3C, D), suggesting a greater capacity to entrain the network activity and modulate intact cortico-cortical connectivity in the rat brain. Despite the greater number of stimulation pulses and the higher mean firing rate observed during the stimulation phase (see Supplementary Materials, Figure S1 A, B), the stimulus-associated activity induced by the RS treatment was lower than ADS and characterized a lower degree of correlation with the stimuli (Figure 3C and D).

In previous awake demonstrations of ADS (Jackson *et al*., 2006; Guggenmos *et al*., 2013), continual pairing of the trigger and target sites led to long-term changes in both neuronal spiking patterns and behavioral outcomes. We attempted to induce these long-term changes with limited (80 minute) stimulation periods daily to more closely mimic what might be applied in a therapeutic setting. Under this paradigm, there was no observable day over day benefit with respect to firing rate. Despite this, daily ADS application over a 21 day period led to increased synaptophysin expression compared to RS, despite receiving significantly less overall stimulation. Synaptophysin, a glycoprotein highly expressed in presynaptic vesicles (Wiedenmann & Franke, 1985), is a marker for synaptic plasticity that has been previously used in stimulation studies (Adkins *et al*., 2008; Gellner *et al*., 2019). Marked increases in synaptophysin are indicative of increased synaptic density which indicates a change in synaptic plasticity, which would be expected if ADS was inducing Hebbian plasticity (Thiele *et al*., 2000). Adkins *et al*. (Adkins *et al*., 2008) found similar increases in synaptic count in sensorimotor cortex following epidural stimulation of the neocortex in an open-loop manner, Cooperrider *et al*. found that chronic contralateral cerebellar deep brain stimulation promotes the perilesional synaptophysin expression in a rodent model of focal ischemia in the cerebral cortex (Cooperrider *et al*., 2014). Synaptogenesis also occurs in late phase motor skill learning (Kleim *et al*., 2004). In all conditions, there was a stable, low level of synaptophysin expression within sections encompassing RFA (the location of the trigger electrode) and referenced baseline synaptophysin expression (Korematsu *et al*., 1993; Wallrafen & Dresbach, 2018). Similar patterns of synaptophysin expression were also found in the hemisphere contralateral to the electrode arrays in both RFA and S1. In contrast, increased fluorescent intensity in S1 of the left hemisphere (see Figure 4)—particularly around the location of electrode placement—suggests that the effect of stimulation is both hemisphere- and region-specific. ADS significantly increased synaptophysin expression in the left S1 compared to RS (see Figure 4B), suggesting that even the short-term neurophysiological effects of ADS described above, might be associated with greater synaptic plasticity than random, open-loop stimulation alone.

Although many experimental protocols have been developed to optimize synaptic potentiation in various model systems, the sign and magnitude of synaptic potentiation are strongly dependent upon the frequency and pattern of stimulation (Feldman, 2009; 2012). Because ADS is dependent on the intrinsic activity within the neural tissue rather than being driven artificially, it may be more optimal for driving synaptic potentiation than other paradigms such as paired-pulse or theta burst stimulation.

Taken together our findings further confirm the ability of Hebbian-inspired protocols to modulate neurophysiological activity by potentiating cortico-cortical connections between distant cortical locations in healthy brains. There is clinical significance in learning that ADS and RS can impact neuroplasticity in the absence of acute brain injury. It is known that synaptogenesis and modulation of synaptic strength through LTP are essential in motor learning and motor recovery following stroke (Wathen *et al*., 2018). Moreover, there is evidence to suggest that post-stroke recovery is associated with the formation of new synapses, as evidenced by animal models demonstrating increased synaptophysin expression in the perilesional cortex (Madinier *et al*., 2013).

Within this experimental framework ADS has shown to be consistently more efficient than RS in both altering single units’ firing rate and evoking stimulus-associated neuronal activity of RFA, therefore proving its greater efficacy to functionally bridge distant neural pathways. In addition, the long-term (21 days) application of ADS rather than RS was able to promote synaptogenesis at the stimulation site (S1), suggesting that structural changes may be induced through artificially bridging neural connections. This further reinforces ADS as a tool for both promoting recovery after brain injury as well as a tool for investigating basic mechanisms of plasticity within and between brain regions.

## Acknowledgements

The authors would like to thank Caleb Dunham from the University of Kansas Medical Center for technical assistance with spike analysis and data management and both Dr Marianna Semprini and Dr Valentina Pasquale from the Istituto Italiano di Tecnologia (IIT) for their useful discussion on computational analysis of the chronic acquisitions.

The authors are especially thankful to Dr Lorenzo De Michieli, Director of the Rehab Technologies Lab, for supporting the research activities conducted by the IIT authors.

## Supplementary Materials

**Figure S1.**
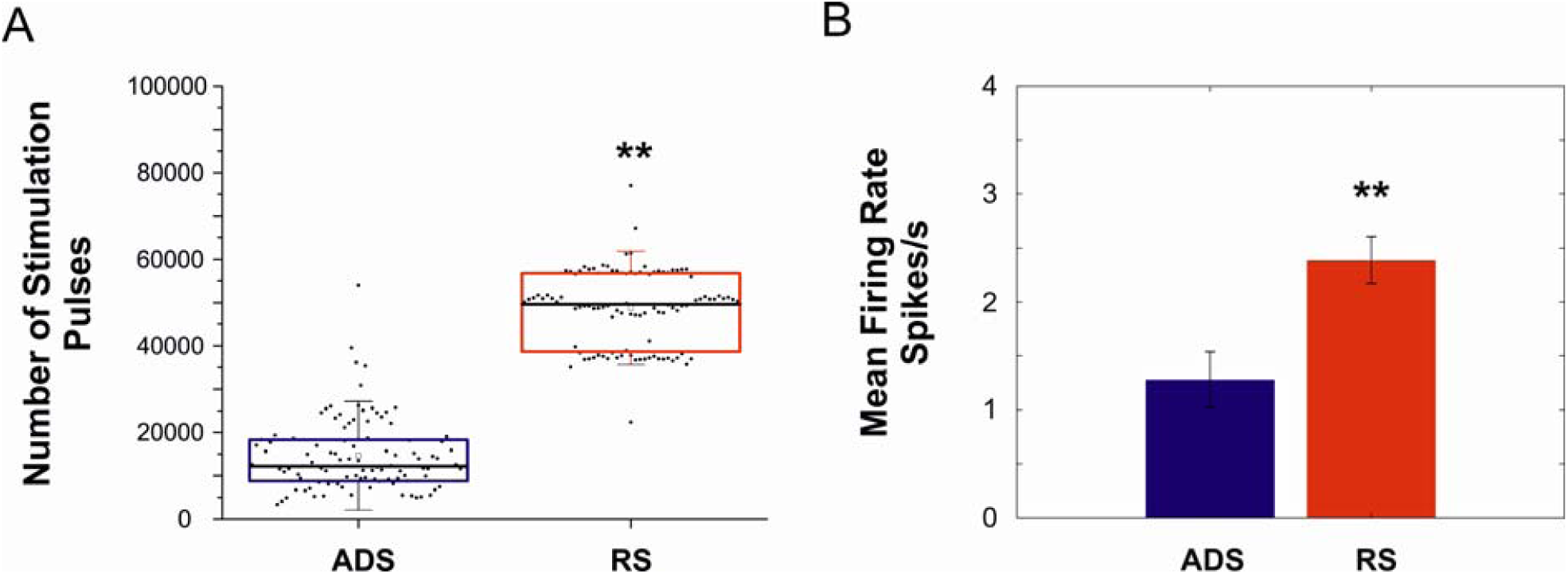
Comparison between ADS and RS. A) Number of pulses provided by the two stimulation protocols, ADS (blue) and RS (red). B) Mean Firing Rate (median±SE of median) calculated during the stimulation phase (Stim) for the two groups. Data pooled across subjects and days for each group. (**p < 0.001; Mann-Whitney Rank Sum test). For each box plot (A), the small central square indicates the mean, the central line illustrates the median and the box limits indicate the 25th and 75th percentiles. Whiskers represent the 5th and the 95th percentiles.

**Figure S2.**
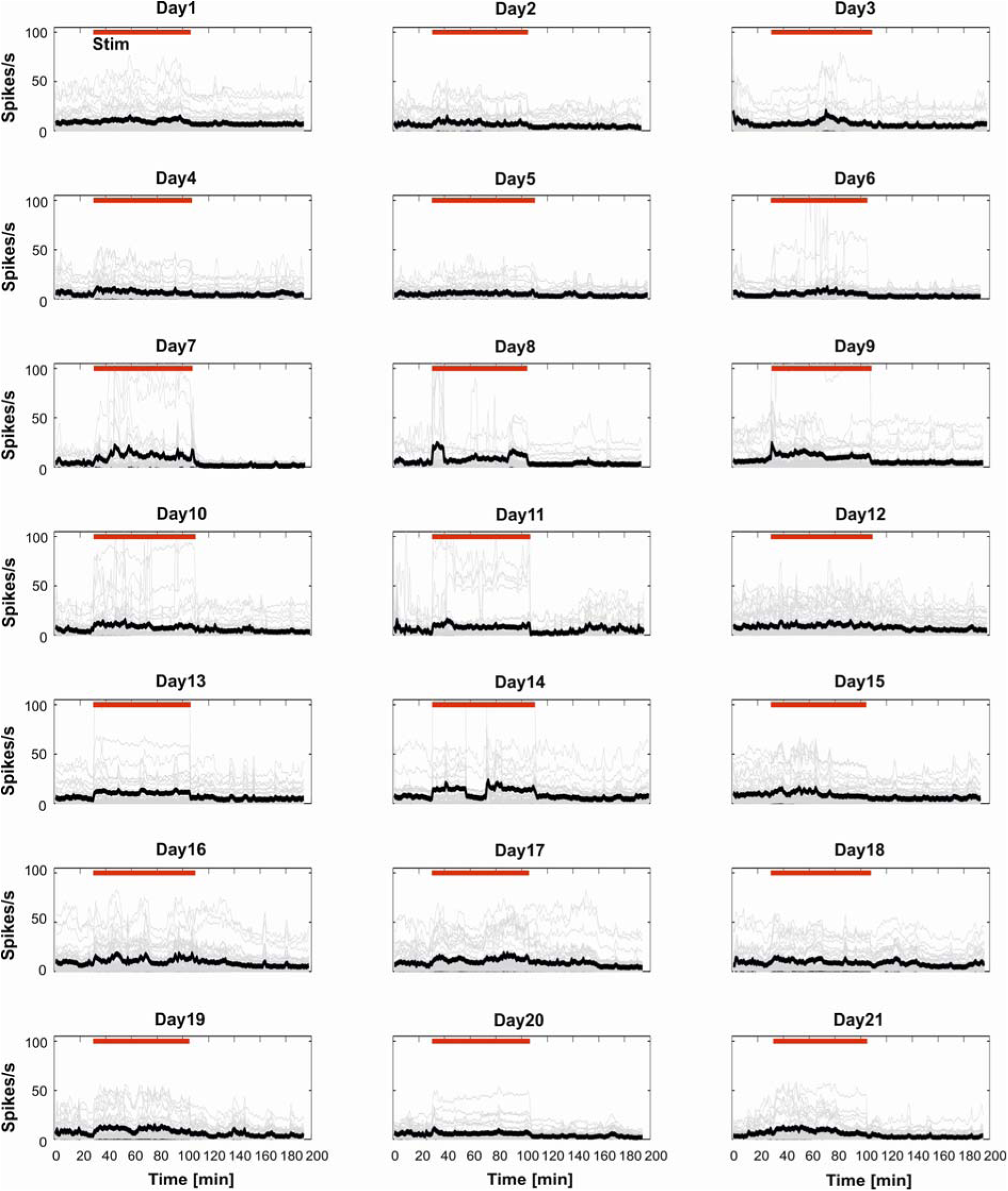
Firing rate does not change across days. Each box represents the instantaneous firing rate (IFR) in 1 min bin of all single units (thin gray lines) and the average IFR (thick black line) calculated during each of the 21 days of recordings over the entire duration of a representative experiment. Red bar indicates the Stim phase.

**Figure S3.**
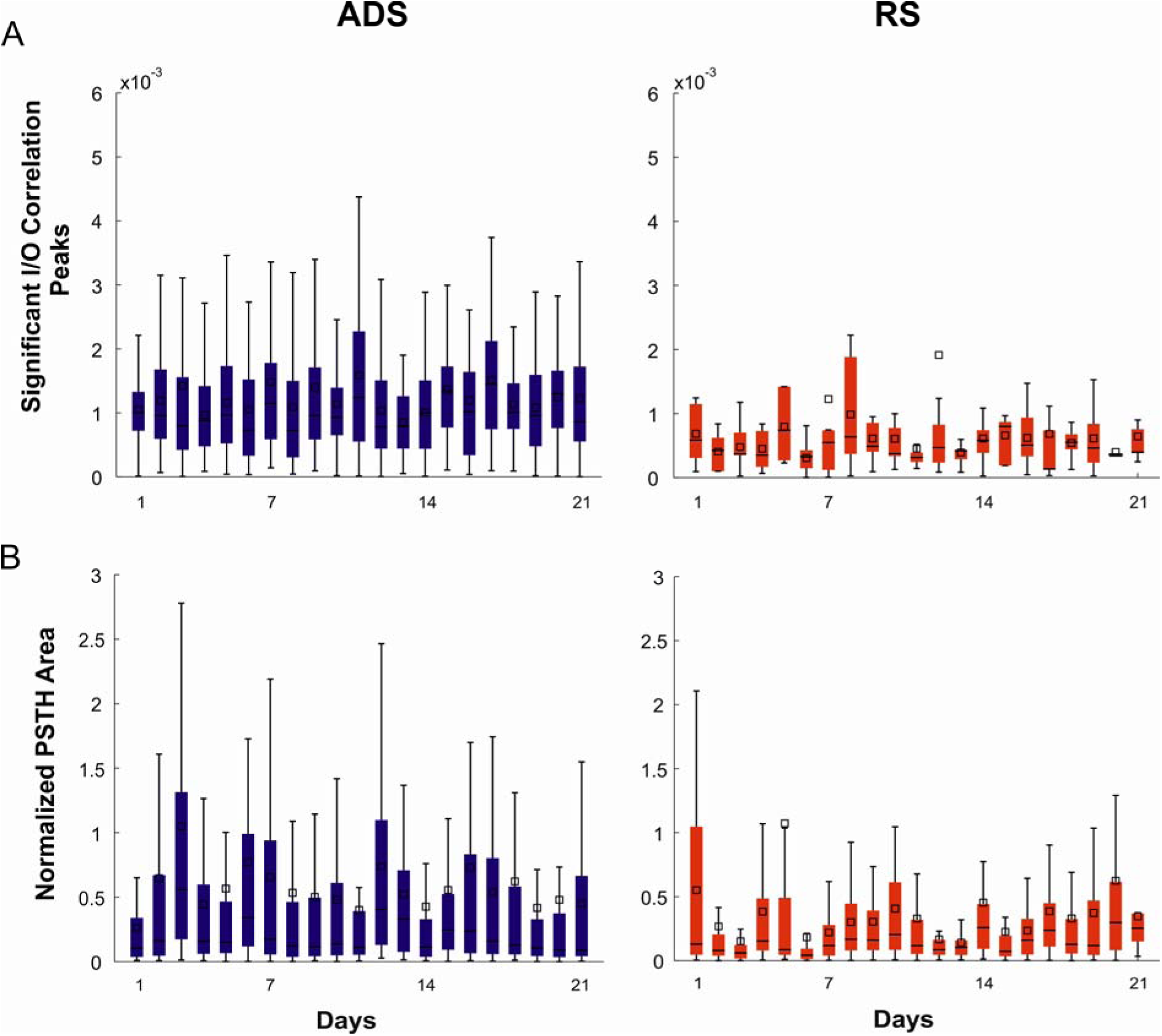
Correlation and PSTH across days. A) I/O Correlation values across days for the two stimulation groups. B) Normalized PSTH areas across days for ADS and RS groups.

